# Spontaneously emerging patterns in human motor cortex code for somatotopic specific movements

**DOI:** 10.1101/2025.04.26.650747

**Authors:** Lu Zhang, Lorenzo Pini, Gordon L Shulman, Maurizio Corbetta

## Abstract

The role of spontaneous brain activity remains a key question in neuroscience. While prior work shows sensory regions preferentially replay stimulus-specific patterns (e.g., faces in fusiform face area), we investigated whether motor cortex replays reflect somatotopic organization. Using fMRI, we compared resting-state activity to task-evoked patterns during specific movements (finger, toe, tongue). Spontaneous activity in each effector-specific motor subregion (hand, foot, mouth) exhibited greater similarity to task patterns of its corresponding movement than to non-preferred ones. For instance, hand movement patterns replayed more in hand regions than elsewhere. Positive rest-task correlations emerged for preferred movements, while negative correlations characterized non-preferred ones (except toe). This structured somatotopic organization suggests spontaneous activity encodes functional specialization, not just activation strength, mirroring task-evoked representations.

## Introduction

A fundamental challenge in modern neuroscience is understanding how the brain’s spontaneous activity and dynamics influence perception, movement, memory, and thought. Even in the absence of sensory input or task engagement, the brain remains constantly active, generating complex dynamical patterns across cortical and subcortical regions [1–5].

The function of spontaneous activity remains obscure. Mainstream models of cortical processing model spontaneous activity as random noise fluctuations that facilitate feedforward postsynaptic activation, modulated by endogenous signals like attention or reward [6–9]. However, it is now well established that spontaneous activity is correlated in space and time in distributed cortical-subcortical networks.

Different hypotheses have been proposed to account for this organization. Large-scale networks may represent emergent properties of anatomical circuitries driven by noise [10, 11]. Alternatively, they may underlie off-line homeostatic or learning mechanisms related to the history of task activation [12–14], which function as a scaffold during task activation [3, 13–17]. Recently, we [18] have proposed that spontaneous activity replays are priors of generative models that encode the most common repertoire of sensory, motor, and cognitive variables of individual behaviour and that these priors are embedded in patterns of spontaneous activity (see also Wittkuhn, Chien [19] for a similar view). One of the predictions of this theory is that priors (replays) shall be specific for the variable that they are coding for.

There is growing experimental evidence for this ‘representational’ hypothesis. First, at rest, in the absence of any task or stimulation, spatial patterns of activity coding for specific visual stimuli (e.g. faces) spontaneously emerge (replay) in human visual cortex. Interestingly, these spontaneous replays occur more commonly for natural than synthetic visual stimuli [20–22]. Second, replays also occur for movement patterns, especially movements that are ecological and frequent [23]. They have been recorded in human motor cortex [24], fronto-parietal association cortex [25], and visual cortex [26]. Moreover, in human somatosensory cortex, patterns related to hand visual stimuli replay more frequently than patterns for non-hand visual stimuli [27]. Similar results have been described across species [28]. In mice, a limited number of spontaneous spatiotemporal patterns recapitulate and explain the majority of task-related patterns [29].

Most of the studies described above have looked for spontaneous activity replays comparing natural vs. synthetic stimuli (e.g., [21, 22, 27, 30]) or ecological vs. non-ecological movements (e.g., [24, 25]). However, the specificity of these patterns for a specific cortical region or system has been less studied. Only Kim et al. [22] showed that category-specific replays (e.g. face or scene patterns) occur more frequently in the corresponding category-specific area (e.g., face replays in fusiform face area vs. scene replays in the parahippocampal place area).

In this study we examined the specificity of motor-related replays in the human motor cortex, taking advantage of its somatotopic organization. Based on the “representation” theory, in the case of movements, they shall respect the somatotopic specificity typical of motor control in the cortex. The human motor cortex is organized in sub-regions along the precentral gyrus that represent different effectors (hand, foot, mouth)[31]. Different subregions and corresponding fronto-parietal loops also represent different ecological actions [32]. Recently, Gordon and colleagues challenged the canonical homuncular organization of M1 reporting that effector-specific regions (foot, hand, mouth) are separated by inter-effector regions that are putatively important for the integration of movements and are preferentially connected with premotor regions [33].

To test the effector specificity of movement-related replays we first measured multivariate patterns of task activation for movement of different effectors, specifically finger tapping, toes squeezing, and tongue moving. These patterns were then compared (spatial correlation) with spontaneously occurring multivariate patterns of activity during resting state scans. This comparison was separately carried out for different subregions of the precentral gyrus (foot, hand, mouth) taken from the recent atlas of [33].

Based on these findings, we extended previous observations of the representational pattern within the motor cortex to assess whether rest-task similarity is observed within these effector-specific regions or whether rest-task “replay” emerges as a continuous property of the whole motor cortex. Specifically, we examined if spontaneously emerging activity patterns in effector-specific (foot, hand, and mouth) areas [33] represent movement-specific patterns during rest. In line with the representation hypothesis of spontaneous activity, we made specific predictions: spontaneous multivoxel activity patterns, i.e., patterns of activity observed at rest in the absence of any stimulation, in functionally specialized motor regions would be more related to the activity patterns evoked by a matched movement (e.g., hand region for hand movements) than by non-matched ones. This predicted relationship between spontaneous and task-evoked multi-voxel patterns within motor effector-specific regions would reinforce the assumptions of entrainment of task-evoked patterns into spontaneous activity.

## Results

### Similarity between spontaneous and task-evoked activity patterns

The effector-specific regions of interest (ROIs) from [33] and the workflow of the analysis are shown in **Fig. 1**. In each participant, we measured multi-voxel patterns of activation in different effector-specific ROIs (hand, foot, mouth) from the atlas of Gordon et al. [33] during finger tapping, toe squeezing, and tongue moving, respectively (see method paragraph). These multi-voxel patterns representing the mean spatial activation pattern for each ROI for each movement were then spatially correlated with similar multi-voxel patterns spontaneously emerging frame-by-frame during the resting state measured prior to the movement. The frequency distribution of spatial correlation values (Pearson’s r) for task-to-rest (task:rest) patterns were computed for each participant, movement, and ROI. As a dependent variable, as in previous work [22], we identified the 90^th^ percentile of the r-value distribution (referred to as U90). U90 values for each participant, movement, and ROI were entered in a within-subject repeated measures analysis of variance (ANOVA).

**Figure 1.**
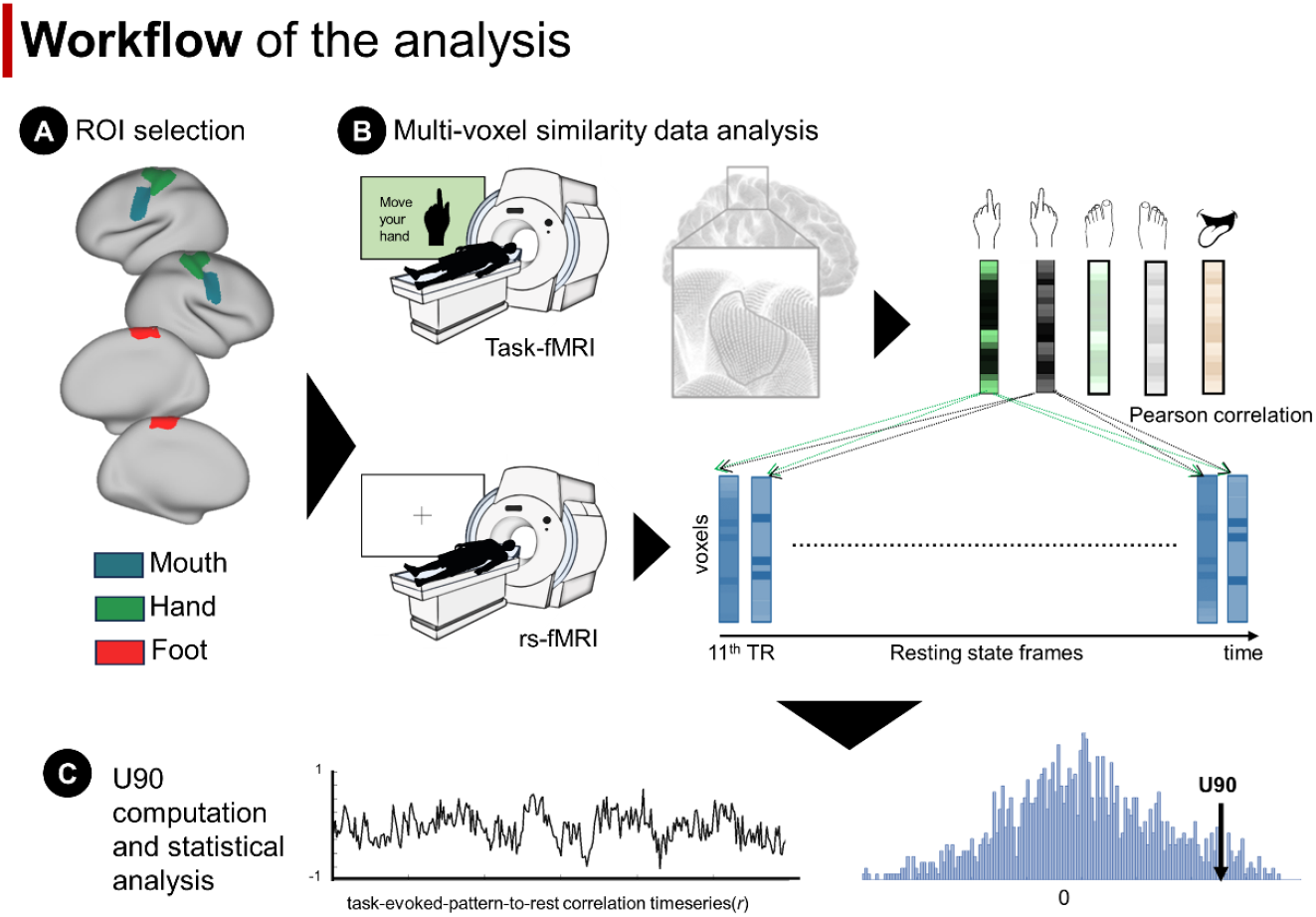
Flow chart of the data analysis. A) Regions of interest (ROIs) based on the atlas from [33]. B) The activation patterns evoked by each motor task were computed using a general linear model (GLM). This procedure yielded five vectors, one for each motor task. A similar voxel vector procedure was adopted for resting-state data leading to multiple vectors, one for each TR. Task-evoked patterns were correlated with resting-state data using Pearson correlation. C) To compare the strength of the correlation, the upper 90% of the distribution (U90) values were computed.

In the primary analysis that includes all the participants, we found a significant interaction effect between movement patterns (five movements considered: tongue moving, left/right fingers tapping and left/right toes squeezing), hemisphere (left, right), and ROI (hand, foot, mouth) (*F*[8,792]=21.73, *p*<0.001, *η*_*p*_^2^=0.180)(**Table 1**). The two-way interaction between movement patterns and ROI computed separately for the left and right hemispheres showed significant effects for both hemispheres (left: *F*[8,792]=28.87, *p*<0.001, *η*_*p*_^2^=0.226; right: *F*[8,792]=26.21, *p*<0.001, *η*_*p*_^*2*^=0.209)(**Table 1**). Further, the two-way interaction between movement patterns and hemisphere computed separately for hand, foot, and mouth ROIs showed significant effects (hand ROI: *F*[4,396]=19.67, *p*<0.001, *η*_*p*_^*2*^=0.166; foot ROI: *F*[4,396]=10.60, *p*<0.001, *η*_*p*_^*2*^=0.097; mouth ROI: *F*[4,396]=18.81, *p*<0.001, *η*_*p*_^*2*^=0.160). These ANOVAs indicate that task:rest spatial correlation activity patterns in human motor cortex were significantly different for different movements across different ROIs in the two hemispheres. To work out the effect of different movements in different ROIs, we first ran a two-way ANOVA with ROI (hand, foot, mouth) and hemisphere (left, right) as factors computed separately for different movement patterns. There was a significant interaction ROI x hemisphere for hand and foot patterns (left foot pattern: *F*[2,198]=16.55, *p*<0.001, *η*_*p*_^*2*^=0.143; right foot pattern: *F*[2,198]=11.98, *p*<0.001, *η*_*p*_^*2*^=0.108; left hand pattern: *F*[2,198]=3.80, *p*=0.024, *η*_*p*_^*2*^=0.037; right hand pattern: *F*[2,198]=34.88, *p*<0.001, *η*_*p*_^*2*^=0.261), while a marginally significant effect was found for the tongue pattern (*F*[2,198]=3.05, *p*=0.049, *η*_*p*_^*2*^=0.030). These results were essentially replicated when the main dataset was split into two different sub-datasets (n=50) (**Table S1**).

**Table 1.** Interaction analysis for all the participants.

|  | All results |  |  |
| --- | --- | --- | --- |
| | *F*-statistic | *p*-value | $\eta_p^2$ |
| Three-way Interactions: Movement patterns x Hemisphere x ROI | 21.73 | <0.001 | 0.180 |
| Two-way Interactions |  |  |  |
| For each hemisphere: Movement patterns x ROI |  |  |  |
| Left hemisphere | 28.87 | <0.001 | 0.226 |
| Right hemisphere | 26.21 | <0.001 | 0.209 |
| For each ROI: Movement patterns x Hemisphere |  |  |  |
| Hand ROI | 19.67 | <0.001 | 0.166 |
| Foot ROI | 10.60 | <0.001 | 0.097 |
| Mouth ROI | 18.81 | <0.001 | 0.160 |
| For each movement patterns: Hemisphere x ROI |  |  |  |
| Left foot patterns | 16.55 | <0.001 | 0.143 |
| Left hand patterns | 3.80 | 0.024 | 0.037 |
| Right foot patterns | 11.98 | <0.001 | 0.108 |
| Right hand patterns | 34.88 | <0.001 | 0.261 |
| Tongue patterns | 3.05 | 0.049 | 0.030 |

Finally, to understand the association between spontaneous and movement-evoked patterns within each movement-specific region, we ran a one-way ANOVA with movement as a factor (tongue moving, left/right fingers tapping and toes squeezing) for each ROI, separately. There was a significant modulation of movement in the hand and mouth ROIs (left-hemisphere hand ROI: *F*[4,396]=26.63, *p*<0.001, *η*_*p*_^*2*^=0.212; right-hemisphere hand ROI: *F*[4,396]=27.85, *p*<0.001, *η*_*p*_^*2*^=0.220; left-hemisphere mouth ROI: *F*[4,396]=39.71, *p*<0.001, *η*_*p*_^*2*^=0.286; right-hemisphere mouth ROI: *F*[4,396]=38.69, *p*<0.001, *η*_*p*_^*2*^=0.281) (**Fig. 2** and **Table 2**). Specifically, within the left-hemisphere hand ROI, spontaneous multi-voxel activity patterns were more similar to task activity patterns evoked by right finger tapping (U90=0.130) than toe squeezing (*p*<0.001, left U90=0.114, right U90=0.103), tongue moving (*p*<0.001, U90=0.092), and left finger tapping (*p*<0.001, U90=0.113) after Bonferroni multiple comparison correction **Fig. 2 and Table 2**).

**Table 2.** Statistical results for the main effect of motor patterns within each region of interest (ROI) for all the participants. Effector-specific ROI preferred U90 values are highlighted in bold. If the highest values do not correspond to the effector-specific ROI preference, they are highlighted in italics.

|  | Main effect of movement patterns |  |  |  |  |  |  |  |
| --- | --- | --- | --- | --- | --- | --- | --- | --- |
| | *F* | *p* | $\eta_p^2$ | U90 values | | | | |
|  |  |  |  | Left foot pattern | Left hand pattern | Right foot pattern | Right hand pattern | Tongue pattern |
| Left Hand ROI | 26.63 | < 0.001 | 0.212 | 0.114 | 0.113 | 0.103 | **0.130** | 0.092 |
| Right Hand ROI | 27.85 | < 0.001 | 0.220 | 0.094 | **0.120** | 0.112 | 0.105 | 0.090 |
| Left Foot ROI | 4.51 | 0.001 | 0.044 | 0.105 | *0.122* | **0.119** | 0.115 | 0.118 |
| Right Foot ROI | 8.39 | < 0.001 | 0.078 | **0.118** | 0.115 | 0.103 | 0.121 | *0.127* |
| Left Mouth ROI | 39.71 | < 0.001 | 0.286 | 0.129 | 0.138 | 0.132 | 0.121 | **0.161** |
| Right Mouth ROI | 38.69 | < 0.001 | 0.281 | 0.131 | 0.130 | 0.134 | 0.148 | **0.171** |

**Figure 2.**
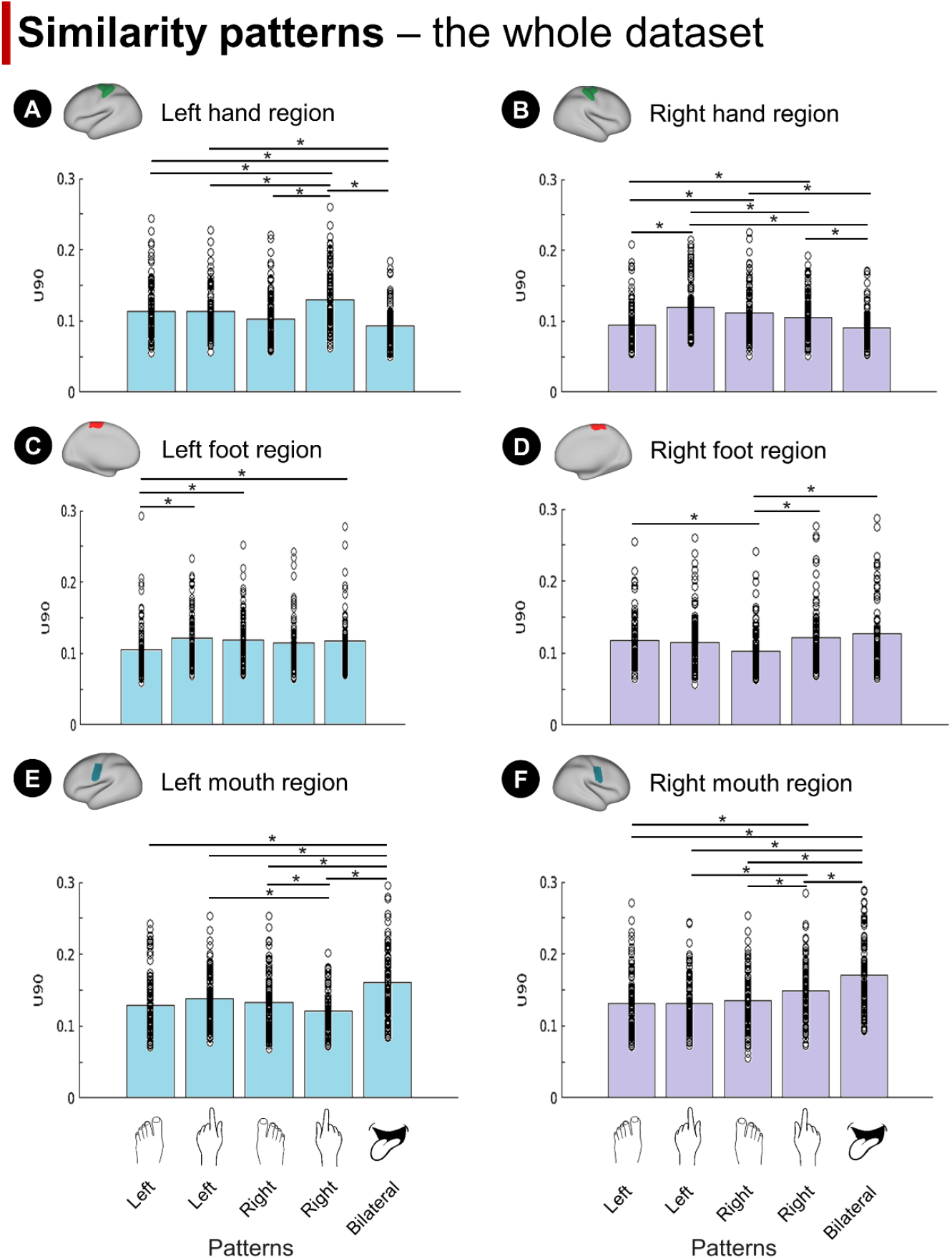
Similarity patterns in the whole dataset. Rest-task similarity patterns for different motor patterns in the effector-specific regions of interest of the motor cortex. Each bar represents the U90 values for each motor pattern within each region. A-B panels: left- and right-hemisphere hand region; C-D panels: left- and right-hemisphere foot region; E-F panels: left- and right-hemisphere mouth region; * marks significant difference corrected with Bonferroni multiple comparison.

Similarly, within the right-hemisphere hand ROI, evoked activity patterns from left finger tapping (U90=0.120) were more similar to spontaneous activity patterns than left toe squeezing (*p*<0.001, U90=0.094), tongue movements (*p*<0.001, U90=0.090), and right finger tapping (*p*<0.001, U90=0.105) after Bonferroni multiple comparison correction. Within the mouth ROIs, regardless of left and right hemisphere, tongue movements (left U90=0.161; right U90=0.171) were more represented in the resting state as compared to finger tapping and toe squeezing patterns (bilateral mouth ROI: tongue pattern-vs-left/right hand/foot pattern: *p*<0.001).

In the left and right foot ROIs, we also found a main effect of movement pattern (left-hemisphere foot ROI: *F*[4,396]=4.51, *p*=0.001, *η*_*p*_^*2*^=0.044; right-hemisphere foot ROI: *F*[4,396]=8.39, *p*<0.001, *η*_*p*_^*2*^=0.078). Post-hoc analyses indicated that resting state patterns in the left hemisphere foot ROI were more like evoked patterns for right than left toes squeezing (*p*=0.011, right U90=0.119, left U90=0.105). In the right hemisphere foot ROI, resting state patterns were more similar for left toe vs. right toe squeezing (*p*=0.009; left U90=0.118, right U90=0.103)(**Fig. 2 and Table 2**). The results for the remaining movements are reported in **Supplementary Materials**. These results were mostly echoed in both replicative sub-datasets, as highlighted in **Fig. S1, Fig.S2**, and **Table S2**.

In summary, these findings show in the whole dataset of participants, as well as in sub-sets of participants, that multivariate patterns of activity emerge in human motor cortex in the resting state that resemble patterns of evoked activity for different movements. Interestingly, the frequency of these patterns at rest varies somatotopically, i.e., they occur more frequently for patterns that are specific for the effector-specific subregion of the motor strip. For instance, finger-tapping movement patterns occur more frequently than foot patterns in the hand sub-region, and so on. Moreover, the similarity between rest and movement patterns is also hemisphere specific, i.e., left finger tapping patterns occur more frequently in the right-hand than left-hand motor ROI, and so on for other movements.

*Positive and negative correlations between U90 values and activation values* The representation hypothesis [18] maintains that task-evoked patterns influence and synchronize with spontaneous activity patterns during development and through experience. Thus, we tested the relationship across participants between the magnitude of the movement-evoked response and rest-task similarity (i.e., the similarity between movement-evoked patterns and spontaneous activity patterns as indexed by the U90 value). U90 values showed a positive correlation with the activation strength for preferred movement patterns in movement-specific regions (e.g., left-hemisphere hand ROI) (**Fig. 3**). The greater the activation magnitude of a movement, the higher the U90 value. Specifically, in the left-hemisphere hand ROI, U90 values for right finger tapping patterns were positively correlated with activation values evoked by right finger tapping (*r*=0.24, *p*=0.015); similarly, in the right-hemisphere hand ROI, U90 values for left finger tapping patterns were positively correlated with activation values evoked by left finger tapping (*r*=0.27, *p*=0.007); in the bilateral mouth ROI, we also observed a significant positive correlation between U90 values and activation values for tongue patterns (left-hemisphere mouth ROI: *r*=0.26, *p*=0.008; right-hemisphere mouth ROI: *r*=0.21, *p*=0.037).

**Figure 3.**
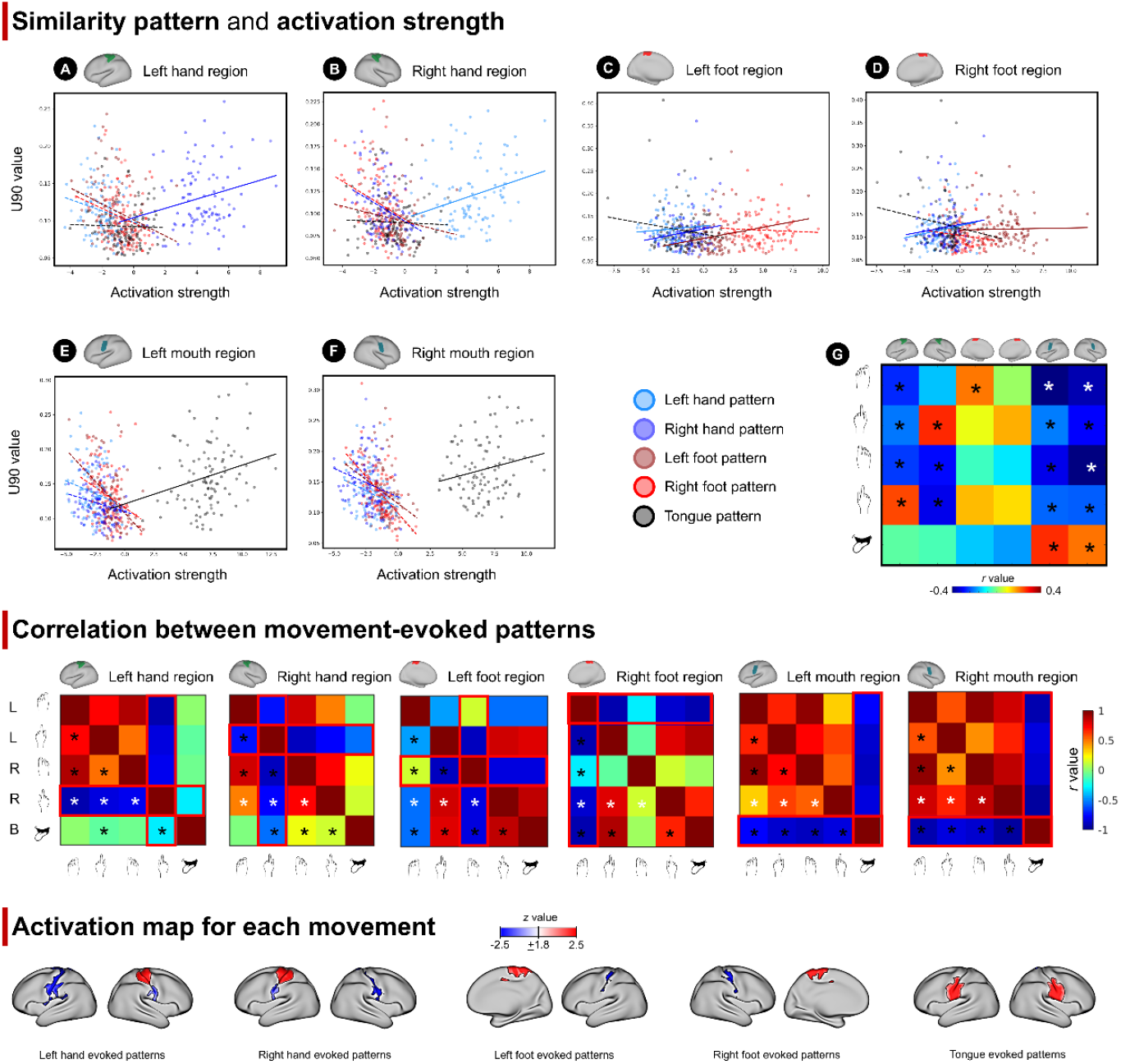
Relationship between U90 and activation strength within motor-specific regions for all the participants. Top panel: A) Left-hemisphere hand region; solid lines show positive correlations while dashed lines show negative correlations; B) Right-hemisphere hand region; C) Left-hemisphere foot region; D) Right-hemisphere foot region; E) Left-hemisphere mouth region; F) Right-hemisphere mouth region; G) The correlation values between U90 and activation strength in each ROI for each movement. * marks significant correlation. **Middle panel:** Correlation matrix of movement-evoked patterns. The red border highlights the correlations between preferred and non-preferred patterns. These correlation matrices are symmetric. Significance is marked only in the lower triangle area. L, left; R, right; B, bilateral. **Bottom panel:** Activation map for each movement evoked patterns.

Interestingly, we also observed a consistent negative correlation: U90 values were negatively correlated with activation values for non-preferred movement patterns in movement-specific regions. Specifically, in the left-hemisphere hand ROI, U90 values of task:rest pattern correlation for non-preferred movement patterns (to the ROI of interest) were negatively correlated with activation values evoked by the same movements (left toe squeezing: *r*=-0.27, *p*=0.007; left finger tapping: *r*=-0.21, *p*=0.038; right toe squeezing: *r*=-0.26, *p*=0.010); likewise, in the right-hemisphere hand ROI (right toe squeezing: *r*=-0.29, *p*=0.003; right finger tapping: *r*=-0.32, *p*=0.001). Additionally, in the bilateral mouth ROI, U90 values for non-preferred movement patterns were negatively correlated with activation values evoked by the same movements (left-hemisphere mouth ROI: left toe squeezing: *r*=-0.47, *p*<0.001; right toe squeezing: *r*=-0.32, *p*=0.001; left finger tapping: *r*=-0.20, *p*=0.043; right finger tapping: *r*=-0.22, *p*=0.027; right-hemisphere mouth ROI: left toe squeezing: *r*=-0.36, *p*<0.001; left finger tapping: *r*=-0.30, *p*=0.002; right toe squeezing: *r*=-0.51, *p*<0.001; right finger tapping: *r*=-0.22, *p*=0.023).

To further investigate the significant negative effects observed, we conducted a post-hoc correlation analysis for each region of interest (ROI), calculating correlations across different movement-evoked patterns within each ROI. This analysis revealed overall negative correlations between preferred and non-preferred patterns. Movement-evoked patterns was averaged across participants and then the correlation between averaged patterns were computed and the correlation matrix is shown in **Fig. 3**. Specifically, we observed significant negative correlations between right finger tapping evoked patterns and left toe squeezing(*r*=-0.92, *p*<0.001), left finger tapping (*r*=-0.81, *p*<0.001), right toe squeezing(*r*=-0.79, *p*<0.001), tongue moving (*r*=-0.30, *p*<0.001) evoked patterns within the left-hemisphere hand ROI. Similarly, significant negative correlations between left finger tapping evoked patterns and left toe squeezing(*r*=-0.73, *p*<0.001), right toe squeezing(*r*=-0.90, *p*<0.001), right finger tapping (*r*=-0.76, *p*<0.001), tongue moving (*r*=-0.56, *p*<0.001) evoked patterns within the right-hemisphere hand ROI. In the bilateral mouth ROI, significant negative correlations were also reported between tongue moving evoked patterns and all other patterns (*r*<-0.76, *p*<0.001). The whole-brain activation maps of the different movements are also reported in **Fig. 3** for visualization purposes.

### No significant relationship between U90 values and motor performance

U90 values (predictors: hand and foot patterns within the contralateral specific-ROI, and mouth patterns in the bilateral specific-ROI) for both the linear (linear regression) and non-linear (random forest) models explained approximately 10% of the motor performance variance, not surviving statistical significance threshold (*p*<0.05). Results are reported in **Fig. 4**.

**Figure 4.**
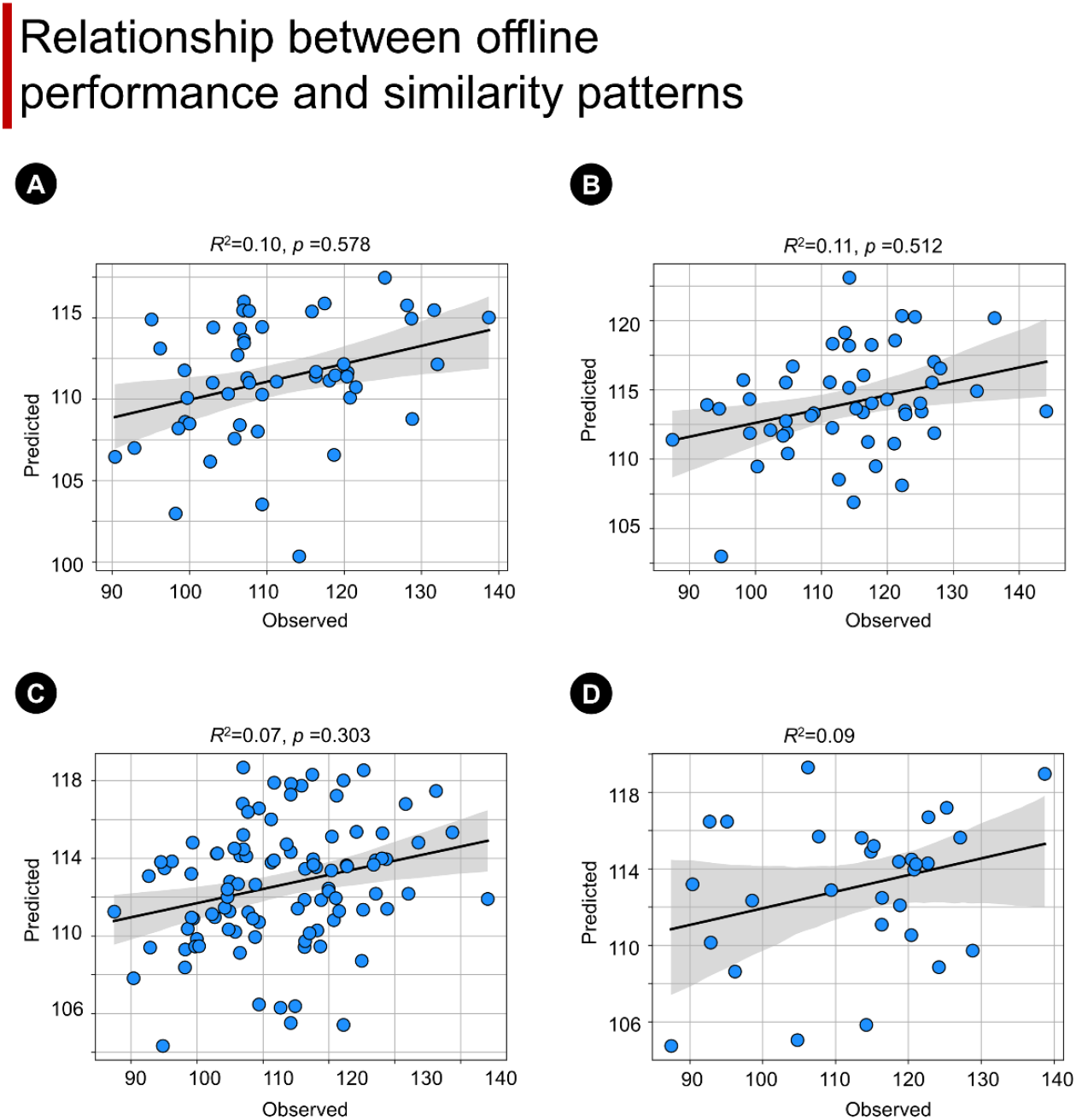
Relationship between U90 and motor task performance. Plots from each panel represent the relationship between predicted and observed behavioral performance of the motor test for both the linear model and random forest model using similarity patterns, i.e., the following U90 values as regressors: left hand and foot pattern within the right ROI; right hand and foot pattern within the left ROI; mouth pattern within bilateral ROI. A) Linear regression for the first replicative sub-dataset. B) Linear regression for the second replicative sub-dataset. C) Linear regression for the pooled dataset including the whole dataset. D) Random forest regression for the pooled dataset. Gray area highlights the confidence interval (CI 95%) of the distribution.

## Discussion

The goal of this study was to assess the representational framework of spontaneous activity in effector-specific regions of the motor cortex by investigating potential links between multivoxel patterns of spontaneous fluctuations and patterns evoked by specific motor movements.

First, we reported that resting state multivoxel activity patterns in effector-specific regions were more similar to the patterns evoked by the regions’ preferred movement. These results suggest that multivoxel spatial patterns of activity at rest show stronger correlations with the multivoxel patterns evoked by tasks or conditions that are commonly encountered or performed or are more congruent with a region’s function, in line with previous findings within the motor cortex [24], visual regions [22], and front-parietal networks [25]. In the present study, the comparison was made between movements involving preferred vs. non-preferred effectors in each effector-specific region. We observed that in the left-hemisphere hand ROI, right finger tapping evoked patterns were more represented in the resting state, while in the hand ROI of the right hemisphere, the left finger tapping evoked patterns were more represented, in line with our hypothesis. Additionally, we observed a similar pattern between spontaneous fluctuations and tongue-evoked activity within the bilateral mouth regions, expanding previous findings to non-limb movements. While for hand movements, the rest-task similarity occurred specifically in the motor cortex contralateral to the performing hand, no lateralization effects were reported for the tongue movement. These results suggest that behavioral representations in the resting state may involve more complex factors than the mapping of effector-specific movements to contralateral effector-specific regions, including motor synergies for actions that involve multiple effectors such as “hand-hand coordination”, “hand-foot coordination”, “bi-foot coordination” and “eye-hand coordination”. Thus, complex movements (e.g., walking and typing) performed during daily life that serve ecological and behaviorally useful functions may be represented in the resting brain [34, 35]. Overall, the representation of task performance during the resting state might play a role in the formation of effector-specific functional zones and their related structural connections during development.

In the foot region, we did not observe significant representation effects. Patterns evoked by toe squeezing were not more represented in the resting state than all other movement-evoked patterns in the bilateral foot region. This null effect might be linked, at least partially, with the movement itself. Toe squeezing is not an ecological/daily movement, and thus, in line with our earlier results, this movement is not expected to be well represented in the resting state [24, 25]. Further studies should assess whether different foot movements (ecological vs uncommon) show the same similarity pattern observed with hands and mouth movements, or whether lower limb movement follows different trajectories, potentially linked with the ontogenesis of these movements [36].

Interestingly, while we observed expected positive relationships between activation strength and rest-task similarity (expressed as U90 values) for region-preferred movement-evoked patterns, we reported a negative relationship for region-unpreferred movement-evoked patterns. These results may reflect how information can be coded in the brain. Computational models indicate that both increases and decreases in activity can be relevant for coding information. Some information states may be explicitly expressed, resulting in a positive match, while others, despite representing the same information, may be latent or dormant, leading to a negative match [37]. These results demonstrate that U90 values do not simply scale positively with the magnitude of task-evoked activity. Instead, they critically depend on how the evoked activity aligns with a region’s functional specialization. Specifically, greater task-evoked activity in a region reduces U90 values when the task involves non-preferred movements for that region. A potential explanation for this effect could involve the role of deactivation during replay. While the prevailing assumption links spontaneous and task-induced signaling to positive task activation, our findings suggest that synchronous deactivation within a region may also play a role in encoding non-preferred movements. Specifically, a low frequency of spontaneous replay for non-relevant movements (e.g., foot patterns in hand sub-regions) may suppress the emergence of strong activation patterns for the same non-relevant movement. Thus, the complex interplay between spontaneous and task-evoked activity is not solely governed by activation phases but also by the deactivation of irrelevant stimuli. This highlights a dual mechanism in which both activation and deactivation contribute to information coding and replay, further suggests a potential mechanism of inhibitory network engagement, where neural representations of non-relevant movements may be actively suppressed during rest to prioritize functionally specialized circuits, implicating dynamic inhibitory processes in maintaining functional segregation within the motor cortex. Future studies should further explore this principle to deepen our understanding of these dynamics.

Finally, we did not observe a significant relationship between motor performance and task-rest similarity. These results are inconsistent with a previous study showing that higher intelligence is related to higher similarity between task and resting brain networks. Specifically, Schultz and Cole [38] compared functional connectivity (FC) during rest and multiple highly distinct tasks and found that the higher the similarity between task and rest FC for a participant, the better the participant’s behavioral performance. However, the null effect (both linear and non-linear) observed in our study could simply reflect the task used to evaluate the association (the 9-hole peg test), since in young healthy participants ceiling effects can mask potential relationships between performance and similarity. Further studies should evaluate whether the degree of similarity is linked with offline behavioral performance in multiple domains. This study has both strengths and limitations. One strength is that the main results from the full sample were largely replicated when the cohort was split into smaller replication sub-datasets [39]. Further, we extended the task-rest similarity representation framework to a new paradigm, the effector-specific regions of the motor cortex. Our results might help to understand the formation of motor functional zones or related structural connections during development. The main limitation of this study is the small set of movements that were included. We cannot interpret the differences between the five movements in terms of familiarity and ecological relevance, as they involve different body parts. Although based on everyday experience, toe squeezing is uncommon relative to walking, whereas tongue movements occur frequently as people talk. Further studies using a spectrum of movements from uncommon to ecological that involve the same body parts, would be informative. Despite this limitation, the primary result of the study, namely the presence of systematic differences in the rest-task similarity of distributed activity patterns in effector-specific regions for different movements was well supported.

## Materials and Methods

### Participants

We included 100 healthy participants (56 females, age range = 22-35 years). from the Human Connectome Project (HCP), WU-Minn Consortium (principal investigators: D. V. Essen and K. Ugurbil; 1U54MH091657) funded by the 16 NIH Institutes and Centers that support the NIH Blueprint for Neuroscience Research and by the McDonnell Center for Systems Neuroscience at Washington University. First, we ran the analysis on the whole cohort of n=100 participants to maximize the precision and accuracy of our results. Then, data were also split randomly into two datasets (n=50 participants each) to replicate results in smaller samples.

### MRI acquisition and task fMRI experimental design

MRI data were acquired using a Siemens 3-T MR system with a 32-channel RF head coil. Resting-state and task functional images were obtained using the same sequence (Gradient-echo EPI sequence): whole-brain echo-planar images with repetition time (TR) = 720 ms, echo time (TE) = 33.1 ms, voxel size = 2 mm^3^ isotropic, 72 interleaved slices. For each participant, high-resolution brain structural images (TR = 2,400 ms; TE = 2.14 ms; voxel size=1 mm3) were acquired using a T1-weighted multi-echo magnetization prepared rapid gradient echo (MPRAGE) sequence.

The imaging procedure was described in Barch, Burgess [40]. During fMRI scanning, participants were asked to perform different movements with hand, foot, and tongue. Specifically, within a block design, participants were asked to tap their left or right fingers, squeeze their left or right toes, or move their tongues according to visual cues. Each movement block lasted 12 s (10 movements) and was preceded by a cue lasting 3 s. In each run, there were 13 blocks: two for tongue, four for hand (balanced for right and left), four for foot (balanced for right and left), and three 15 s fixation blocks. A fifteen-minute resting-state fMRI scan was acquired for each participant. During resting state scans, participants were instructed to relax and keep their eyes open focusing on a projected bright cross on a dark background.

### Data processing

The following preprocessing steps were performed, including gradient unwarping, motion correction, fieldmap-based EPI distortion correction, brain-boundary-based registration of EPI to structural T1-weighted scan, non-linear (FNIRT) registration into MNI152 space, grand-mean intensity normalization, and spatial smoothing was applied using an unconstrained 3D Gaussian kernel of FWHM = 4 mm [40]. Additionally, we regressed out the global (mean) signal from the temporal activity pattern of each voxel. Finally, we normalized the activity levels in each TR by converting them to z-score values, in line with our previous studies [24, 25].

Contrast maps obtained from HCP processed motor task data were included for the following analysis. The same preprocessing steps described above for resting state data were also applied in the motor task dataset. Activity estimates were computed for the preprocessed functional time series from each run using a general linear model (GLM) implemented in FSL’s FILM (FMRIB’s Improved Linear Model with autocorrelation correction). Predictors were convolved with a double gamma “canonical” hemodynamic response function (HRF) to generate the main model regressors. Five predictors were included in the GLM model — right hand, left hand, right foot, left foot, and tongue. Each predictor covered a duration of 12 seconds. Linear contrasts were computed to estimate activation for each movement type, both compared to baseline or all the other movements. In this study, only linear contrasts from each specific movement compared to all the other movements were selected. See [40] for a full description of the dataset and analysis. The average map for each contrast across participants (movement vs all the other movements) is reported in Fig. 3.

### Rest-task multi-voxel similarity analysis

Effector-specific regions were selected as ROIs based on the atlas from [33]. Specifically, three effector-specific regions in both hemispheres, i.e., the hand, foot, and mouth regions, were selected to test the hypothesis that motor-specific patterns would be more represented in the motor effector-specific regions than other motor patterns.

To investigate the similarity between resting-state and task patterns evoked by different motor movements, we performed a multi-voxel linear analysis, based on our previous publications [22, 24, 25]. In detail, within each ROI, we computed the multi-voxel spatial pattern of activation for the five movements considered (i.e., tongue moving, left/right fingers tapping, and left/right toes squeezing). A total of five vectors were computed, one for each motor movement pattern. The length of these vectors represented the number of voxels for each ROI. A vector of the same length was computed for each resting state frame. We then correlated the task-evoked vector with the resting-state data (after removing the first 10 TRs). This procedure yielded n=1190 Pearson correlation values for each movement. To compare the strength of the correlation between task-evoked and resting-state signals, we computed the distribution of these r values. According to our previous analysis [22], we identified the U90 value (the U90 value: higher values indicate higher similarity between task activation and resting-state spontaneous fluctuations). These U90 values were inserted as a dependent variable in an ANOVA, aimed at assessing the interactions between ROIs, movements, and hemispheres, along with the main effects of these factors. The statistical analysis was run within the whole dataset. We also repeated the analyses on both sub-datasets (n=50 each) to assess the robustness of the findings.

Further, we assessed whether higher task activation for specific movements would drive differences between movements in rest-task similarity. To this aim, for each participant and ROI, we computed an averaged movement activation value and correlated it across participants with their U90 value using Pearson correlation.

### Association between U90 values and motor performance

To test the behavioral relevance of intrinsic connectivity modulation in the motor cortex, we tested its relationship with motor task performance. Specifically, we used the 9-hole peg test scores, which evaluates manual dexterity by measuring the time taken to complete a peg-manipulation task in seconds (lower values reflect better task performance) [41–43]. We included participants’ U90 values for finger tapping/toe squeezing in their contralateral hand/foot ROIs (e.g., left finger tapping in the right-hemisphere hand ROI, and vice versa for the right finger tapping) and tongue moving in the bilateral mouth ROI in this analysis. Both linear and non-linear regression models were run, with U90 values as the independent variable and the 9-hole peg test scores as the dependent variable.

The linear model was run independently for both replicative sub-datasets and within the pooled dataset including individuals from both cohorts. Results were analyzed in terms of variance (r-squared) explained by the U90 values for predicting the motor performance. For the non-linear regression, a random forest algorithm was performed, a non-linear ensemble learning method that builds a multitude of decision trees. To ensure enough data during the validation procedure we used the pooled dataset (n=100), and data were split into test and train data with a 30% and 70% proportion, respectively. The training dataset was used to identify the best set of hyperparameters, that is: estimators (from 200 to 2000 with step of 200), depth (from 1 to 10 with step of 1), sample split (2, 5, and 10), samples leaf (1, 2, and 4), and bootstrap (true or false). The best set of parameters identified with a 5-fold cross-validation on the train set was as follows: max depth=9, min samples leaf=4, min samples split=5, number of interactions=600, and bootstrap=True. The fit of the model (r-squared) was then computed as the outcome measure.

## Data and code Availability

### Resource availability

Lead Contact Further information and requests for resources and reagents should be directed to and will be fulfilled by the Lead Contact, Maurizio Corbetta.

### Materials Availability

This study did not generate new unique reagents.

### Data and Code Availability

The neuroimaging data generated during this study are available at cnda.wustl.edu. All code needed to reproduce our analyses is also available at cnda.wustl.edu. The processed dataset and codes could be made available upon reasonable usage request from the corresponding author along with the permission of the Central Neuroimaging Data Archive (CNDA) Center in Washington University in St. Louis (cnda.wustl.edu).

## Author contributions

L.Z. and L.P. designed research and analyzed data; and L.Z., L.P., G.L.S., and M.C. wrote the paper.

## Reference

1. Raichle, M.E. and M.A. Mintun, Brain work and brain imaging. Annu Rev Neurosci, 2006. 29: p. 449–76.

2. György Buzsáki, M., The brain from inside out. 2019: Oxford University Press.

3. Raichle, M.E., The restless brain. Brain Connect, 2011. 1(1): p. 3–12.

4. Buckner, R.L., F.M. Krienen, and B.T. Yeo, Opportunities and limitations of intrinsic functional connectivity MRI. Nat Neurosci, 2013. 16(7): p. 832–7.

5. Carhart-Harris, R.L., K.J. Friston, and E.L. Barker, REBUS and the Anarchic Brain: Toward a Unified Model of the Brain Action of Psychedelics. Pharmacological Reviews, 2019. 71(3): p. 316–344.

6. Corbetta, M. and G.L. Shulman, Control of goal-directed and stimulus-driven attention in the brain. Nat Rev Neurosci, 2002. 3(3): p. 201–15.

7. Tolhurst, D.J., J.A. Movshon, and A.F. Dean, The statistical reliability of signals in single neurons in cat and monkey visual cortex. Vision Res, 1983. 23(8): p. 775–85.

8. Desimone, R. and J. Duncan, Neural mechanisms of selective visual attention. Annu Rev Neurosci, 1995. 18: p. 193–222.

9. Shadlen, M.N. and W.T. Newsome, The variable discharge of cortical neurons: implications for connectivity, computation, and information coding. J Neurosci, 1998. 18(10): p. 3870–96.

10. Deco, G., et al., Resting-state functional connectivity emerges from structurally and dynamically shaped slow linear fluctuations. J Neurosci, 2013. 33(27): p. 11239–52.

11. Barttfeld, P., et al., Signature of consciousness in the dynamics of resting-state brain activity. Proceedings of the National Academy of Sciences, 2015. 112(3): p. 887–892.

12. Lewis, C.M., et al., Learning sculpts the spontaneous activity of the resting human brain. Proceedings of the National Academy of Sciences, 2009. 106(41): p. 17558–17563.

13. Petersen, S.E. and O. Sporns, Brain Networks and Cognitive Architectures. Neuron, 2015. 88(1): p. 207–19.

14. Harmelech, T. and R. Malach, Neurocognitive biases and the patterns of spontaneous correlations in the human cortex. Trends Cogn Sci, 2013. 17(12): p. 606–15.

15. Shine, J.M., et al., Human cognition involves the dynamic integration of neural activity and neuromodulatory systems. Nat Neurosci, 2019. 22(2): p. 289–296.

16. Fiser, J., et al., Statistically optimal perception and learning: from behavior to neural representations. Trends Cogn Sci, 2010. 14(3): p. 119–30.

17. Chanes, L. and L.F. Barrett, Redefining the Role of Limbic Areas in Cortical Processing. Trends in Cognitive Sciences, 2016. 20(2): p. 96–106.

18. Pezzulo, G., M. Zorzi, and M. Corbetta, The secret life of predictive brains: what’s spontaneous activity for? Trends in Cognitive Sciences, 2021.

19. Wittkuhn, L., et al., Replay in minds and machines. Neurosci Biobehav Rev, 2021. 129: p. 367–388.

20. Smith, S.M., et al., Correspondence of the brain’s functional architecture during activation and rest. Proceedings of the national academy of sciences, 2009. 106(31): p. 13040–13045.

21. Wilf, M., et al., Spontaneously Emerging Patterns in Human Visual Cortex Reflect Responses to Naturalistic Sensory Stimuli. Cereb Cortex, 2017. 27(1): p. 750–763.

22. Kim, D., et al., Spontaneously emerging patterns in human visual cortex and their functional connectivity are linked to the patterns evoked by visual stimuli. J Neurophysiol, 2020. 124(5): p. 1343–1363.

23. Zhang, L., et al., Brain-wide dynamic coactivation states code for hand movements in the resting state. Proc Natl Acad Sci U S A, 2025. 122(11): p. e2415508122.

24. Livne, T., et al., Spontaneous activity patterns in human motor cortex replay evoked activity patterns for hand movements. Sci Rep, 2022. 12(1): p. 16867.

25. Zhang, L., et al., Spontaneous activity patterns in human attention networks code for hand movements. J Neurosci, 2023.

26. Stringer, C., et al., Spontaneous behaviors drive multidimensional, brainwide activity. Science, 2019. 364(6437): p. 255.

27. El Rassi, Y., et al., A visual representation of the hand in the resting somatomotor regions of the human brain. Scientific Reports, 2024. 14(1).

28. Dimakou, A., et al., The predictive nature of spontaneous brain activity across scales and species. Neuron, 2025.

29. MacDowell, C.J. and T.J. Buschman, Low-Dimensional Spatiotemporal Dynamics Underlie Cortex-wide Neural Activity. Current Biology, 2020. 30(14): p. 2665–2680.e8.

30. Fiser, J., C. Chiu, and M. Weliky, Small modulation of ongoing cortical dynamics by sensory input during natural vision. Nature, 2004. 431(7008): p. 573–578.

31. Penfield, W. and E. Boldrey, Somatic Motor and Sensory Representation in the Cerebral Cortex of Man as Studied by Electrical Stimulation. Brain, 1937. 60(4): p. 389–443.

32. Graziano, M.S.A., Ethological Action Maps: A Paradigm Shift for the Motor Cortex. Trends Cogn Sci, 2016. 20(2): p. 121–132.

33. Gordon, E.M., et al., A somato-cognitive action network alternates with effector regions in motor cortex. Nature, 2023.

34. Latash, M.L., J.P. Scholz, and G. Schöner, Toward a new theory of motor synergies. Motor control, 2007. 11(3): p. 276–308.

35. Leo, A., et al., A synergy-based hand control is encoded in human motor cortical areas. Elife, 2016. 5.

36. Raffalt, P.C., T. Alkjær, and E.B. Simonsen, Intra- and inter-subject variation in lower limb coordination during countermovement jumps in children and adults. Human Movement Science, 2016. 46: p. 63–77.

37. van Loon, A.M., et al., Current and future goals are represented in opposite patterns in object-selective cortex. Elife, 2018. 7.

38. Schultz, D.H. and M.W. Cole, Higher Intelligence Is Associated with Less Task-Related Brain Network Reconfiguration. J Neurosci, 2016. 36(33): p. 8551–61.

39. Button, K.S., et al., Power failure: why small sample size undermines the reliability of neuroscience. Nature Reviews Neuroscience, 2013. 14(5): p. 365–376.

40. Barch, D.M., et al., Function in the human connectome: task-fMRI and individual differences in behavior. Neuroimage, 2013. 80: p. 169–89.

41. Gershon, R.C., et al., Assessment of neurological and behavioural function: the NIH Toolbox. The Lancet Neurology, 2010. 9(2): p. 138–139.

42. Wang, Y.C., et al., Dexterity as measured with the 9-Hole Peg Test (9-HPT) across the age span. J Hand Ther, 2015. 28(1): p. 53–9; quiz 60.

43. Ruck, L. and P.T. Schoenemann, Handedness measures for the Human Connectome Project: Implications for data analysis. Laterality, 2021. 26(5): p. 584–606.

