## Supplemental materials for "Spontaneously emerging patterns in human motor cortex code for somatotopic specific movements"

**Number of Supplementary Figures: 2**
**Number of Supplementary Tables: 2**

**Supplementary Statistical analysis**

Within the left-hemisphere hand ROI, left finger tapping and left toe squeezing evoked multi-voxel activity patterns showed higher similarity with spontaneous multi-voxel activity patterns than tongue movements (left hand pattern-vs-tongue pattern: Bonferroni corrected $p<0.001$; left foot pattern-vs-tongue pattern: Bonferroni corrected $p<0.001$).

Similarly, within the right-hemisphere hand ROI, right foot and hand patterns showed significantly higher cutoff values than tongue patterns (right foot pattern-vs-tongue pattern: $p<0.001$, right hand pattern-vs-tongue pattern: $p<0.001$) and left foot patterns (right foot pattern-vs- left foot pattern: $p<0.001$, right hand pattern-vs- left foot pattern: $p=0.007$).

We also observed a higher cutoff value for left hand patterns ($p<0.001$) and right foot patterns ($p=0.006$) than right hand patterns in the left-hemisphere mouth ROI, while in the right-hemisphere mouth ROI, differences were also found between right hand patterns and left hand patterns (right hand pattern-vs-left hand pattern: $p<0.001$), and between right hand patterns and left/right foot patterns (right hand pattern-vs-left foot pattern: $p<0.001$; right hand pattern-vs-right foot pattern: $p=0.004$).

In the left foot ROIs, we found that resting state data showed greater similarity to left hand patterns (left hand pattern-vs-left foot pattern: $p=0.001$), tongue moving patterns (tongue moving pattern-vs-left foot pattern: $p=0.028$) than left foot patterns, while in the right-hemisphere foot ROI, resting state data showed greater similarity to tongue patterns and right hand patterns than right foot (tongue pattern-vs-right foot pattern: $p<0.001$; right hand pattern-vs-right foot pattern: $p<0.001$) (**Fig. 2**). These results were mostly echoed in both replicative sub-datasets. See **Fig. S1, Fig.S2,** and **Table S2** for the statistical details.

**Table S1.** Interaction analysis.

| | Replicative dataset 1 | | | Replicative dataset 2 | | |
|---|---|---|---|---|---|---|
| | *F*-statistic | *p*-value | $\eta_p^2$ | *F*-statistic | *p*-value | $\eta_p^2$ |
| Three-way Interactions: Movement patterns x Hemisphere x ROI | | | | | | |
| | 10.39 | <0.001 | 0.175 | 12.13 | <0.001 | 0.198 |
| Two-way Interactions | | | | | | |
| For each hemisphere: Movement patterns x ROI | | | | | | |
| Left hemisphere | 13.89 | <0.001 | 0.221 | 14.85 | <0.001 | 0.233 |
| Right hemisphere | 13.48 | <0.001 | 0.216 | 13.69 | <0.001 | 0.218 |
| For each ROI: Movement patterns x Hemisphere | | | | | | |
| Hand ROI | 8.31 | <0.001 | 0.145 | 11.81 | <0.001 | 0.194 |
| Foot ROI | 3.56 | 0.008 | 0.068 | 9.37 | <0.001 | 0.160 |
| Mouth ROI | 13.16 | <0.001 | 0.212 | 7.87 | <0.001 | 0.138 |
| For each movement patterns: Hemisphere x ROI | | | | | | |
| Left foot patterns | 8.26 | <0.001 | 0.144 | 8.14 | <0.001 | 0.142 |
| Left hand patterns | 5.10 | 0.008 | 0.094 | 2.37 | 0.099 | |
| Right foot patterns | 4.03 | 0.021 | 0.076 | 8.28 | <0.001 | 0.144 |
| Right hand patterns | 15.93 | <0.001 | 0.245 | 19.88 | <0.001 | 0.289 |
| Tongue patterns | 0.93 | 0.398 | | 2.57 | 0.082 | |

**Table S2.** Statistical results for the main effect of motor patterns within each ROI. Effector-specific ROI preferred U90 values are highlighted in bold. If the highest values do not correspond to the effector-specific ROI preference, they are highlighted in italics.

| | Main effect of movement patterns | | | | | | | |
|---|---|---|---|---|---|---|---|---|
| | *F* | *p* | $\eta_p^2$ | U90 values | | | | |
| | | | | Left foot pattern | Left hand pattern | Right foot pattern | Right hand pattern | Tongue pattern |
| | | | Replicative dataset 1 | | | | | |
| Left Hand ROI | 10.60 | < 0.001 | 0.178 | 0.111 | 0.114 | 0.102 | **0.130** | 0.095 |
| Right Hand ROI | 15.65 | < 0.001 | 0.242 | 0.091 | **0.122** | 0.109 | 0.106 | 0.089 |
| Left Foot ROI | 2.81 | 0.027 | 0.054 | 0.104 | *0.122* | **0.115** | 0.115 | 0.119 |
| Right Foot ROI | 4.10 | 0.003 | 0.077 | **0.117** | 0.120 | 0.102 | *0.127* | 0.124 |
| Left Mouth ROI | 24.94 | < 0.001 | 0.337 | 0.126 | 0.139 | 0.131 | 0.120 | **0.162** |
| Right Mouth ROI | 24.05 | < 0.001 | 0.329 | 0.129 | 0.122 | 0.129 | 0.145 | **0.165** |
| | | | Replicative dataset 2 | | | | | |
| Left Hand ROI | 17.14 | < 0.001 | 0.259 | 0.117 | 0.112 | 0.104 | **0.131** | 0.090 |
| Right Hand ROI | 13.05 | < 0.001 | 0.210 | 0.097 | **0.118** | 0.116 | 0.105 | 0.091 |
| Left Foot ROI | 2.21 | 0.069 | | 0.107 | 0.122 | **0.123** | 0.115 | 0.117 |
| Right Foot ROI | 6.38 | < 0.001 | 0.115 | **0.118** | 0.109 | 0.104 | 0.115 | *0.130* |
| Left Mouth ROI | 15.82 | < 0.001 | 0.244 | 0.132 | 0.137 | 0.134 | 0.121 | **0.159** |
| Right Mouth ROI | 16.59 | < 0.001 | 0.253 | 0.134 | 0.138 | 0.140 | 0.151 | **0.176** |

**Figure S1. Similarity patters in the first replicative sub-dataset.** Rest-task similarity patterns for different motor patterns in the effector-specific regions of interest of the motor cortex. Each bar represents the U90 values for each motor pattern within each region. A-B panels: left- and right-hemisphere hand region; C-D panels: left- and right-hemisphere foot region; E-F panels: left- and right-hemisphere mouth region; * marks significant difference corrected with Bonferroni multiple comparison.

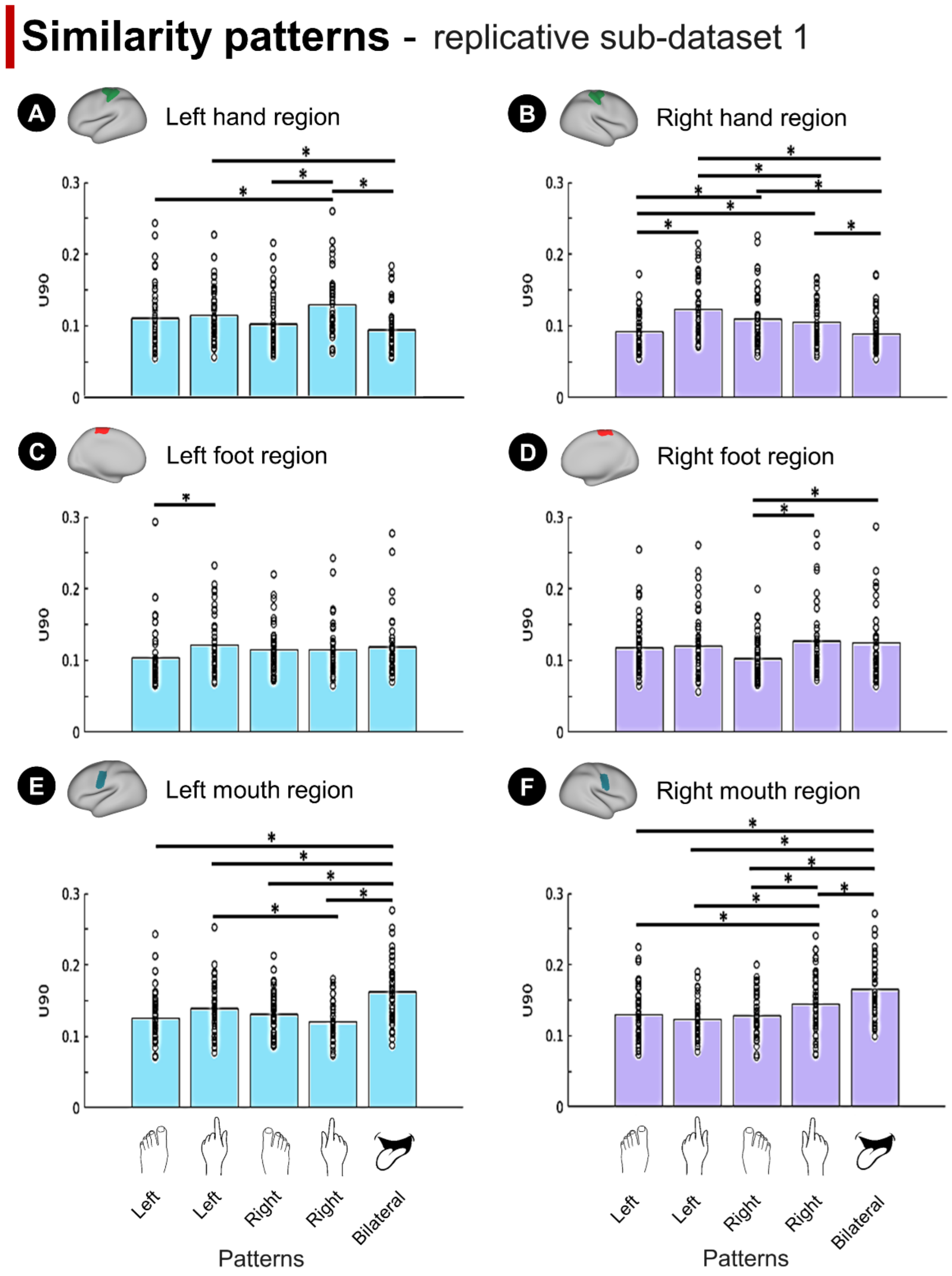

**Figure S2. Similarity patterns in the replicative dataset.**
Rest-task similarity patterns for different motor patterns in the effector-specific regions of interest of the motor cortex. Each bar represents the U90 values for each motor pattern within one region. A-B panels: left- and right-hemisphere hand region; C-D panels: left- and right--hemisphere foot region; E-F panels: left- and right--hemisphere mouth region; * marks significant difference corrected with Bonferroni multiple comparison.

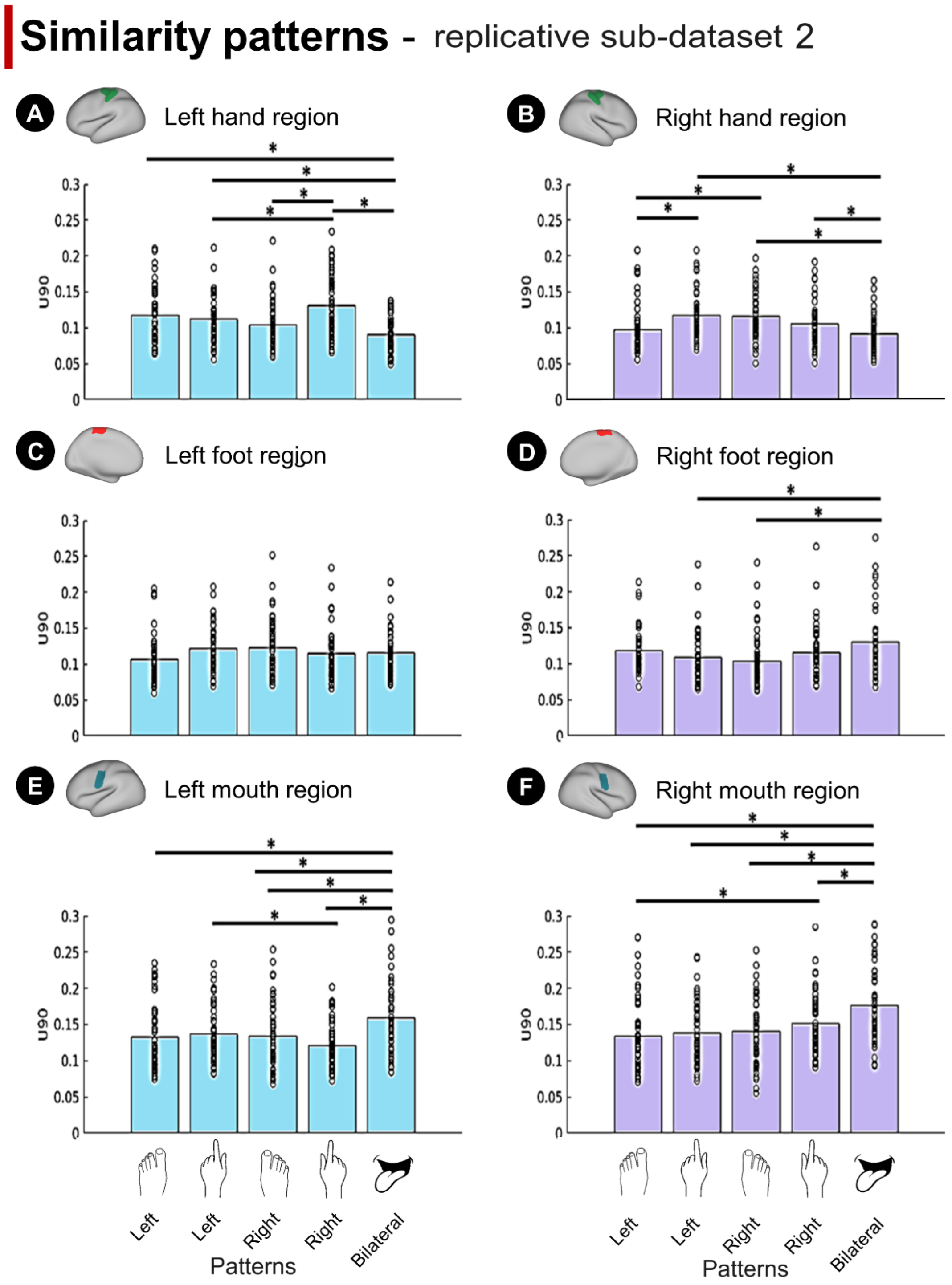
